# Task switching costs contribute to the apparent speeding of multisensory reaction times

**DOI:** 10.1101/2021.03.09.434565

**Authors:** Miriam Albat, Jasmin Hautmann, Christoph Kayser, Josefine Molinski, Soner Ülkü

## Abstract

Faster reaction times for the detection of multisensory compared to unisensory stimuli are considered a hallmark of multisensory integration. While this multisensory redundant signals effect (RSE) has been reproduced many times, it has also been repeatedly criticized as confounding multisensory integration and general task switching effects. When unisensory and multisensory conditions are presented in random order, some trials repeat the same sensory-motor association (e.g. an auditory followed by an auditory trial), while others switch this association (e.g. an auditory followed by a visual trial). This switch may slow down unisensory reaction times and inflate the observed RSE. Following this line of ideas, we used an audio-visual detection task and quantified the RSE for trials derived from pure unisensory blocks and trials from mixed blocks involving a repeat or switch of modalities. The RSE was largest for switch trials and smallest for unisensory trials. In fact, during unisensory blocks the multisensory reaction times did not differ from predictions by the race model, speaking against a genuine multisensory benefit. These results confirm that the observed multisensory RSE can be easily confounded by task switching costs, and suggest that the true benefit of multisensory stimuli for reaction speed may often be overestimated.

## 1. Introduction

Multisensory integration comes with key benefits for perception, such as an improved accuracy or reliability of sensory estimates or a speeding of reaction times (Hershenson, 1962; Molholm et al., 2002; Stein and Rowland, 2020). In fact, when reaction times in a simple detection task are compared between the simultaneously-presented multisensory stimulus and the two unisensory stimuli each presented individually, one usually finds that the multisensory reaction times are substantially faster (Miller, 1982; Schroger and Widmann, 1998; Minakata and Gondan, 2019; Colonius and Diederich, 2020; Shaw et al., 2020). This has been considered a classical hallmark of multisensory perception and is also known as the redundant signals effect (RSE). This multisensory RSE has been reproduced in many variations of the basic experimental setting, and has been compared to neural signatures of multisensory integration (Miniussi et al., 1998; Molholm et al., 2002; Murray et al., 2005). However, a number of studies (Spence et al., 2001; Gondan et al., 2004; Shaw et al., 2020) have suggested that the multisensory RSE obtained in many studies may - at least partly - arise from cognitive effects such as task switching, rather than from a genuine sensory-level benefit of multisensory integration.

The present study was designed to directly revisit the notion that task switching contributes to the observed multisensory RSE. To this end we reproduced the experiment and analyses proposed by Shaw et al. (2020) - see (Gondan et al., 2004) for a similar logic. In that study the RSE was quantified in different experimental contexts and by directly probing for a potential contribution of task switching. The authors capitalized on the notion that the traditional RSE paradigm, in which uni- and multisensory trials are randomly intermingled, effectively comprises both a switch in sensory modality and a switch in response-relevant domains at the same time: a typical task presents the three modality conditions (e.g. auditory, visual and audio-visual) in a pseudo-random order and with variable inter-trial intervals. Hence, participants cannot anticipate the timing or the nature of the upcoming stimulus. However, due to the random order, a given trial may effectively repeat the previous stimulus-response mapping (e.g. an auditory trial followed by an auditory trial) or may induce a switch of this (e.g. an auditory trial followed by a visual trial). When seen in the light of the task-switching literature (Posner et al., 1976; Kiesel et al., 2010; Koch et al., 2010), this either entrails a repeat and a switch of the stimulus-response association. This switch in stimulus-response domains may confound the mere effect of changing the sensory modality when comparing reaction times between conditions (Spence et al., 2001; Gondan et al., 2004; Shaw et al., 2020).

To test whether task switching contributes to the typically observed multisensory RSE, some studies measured the RSE obtained when presenting the two unisensory and the multisensory condition in three separate unisensory blocks, and they measured the RSE obtained in blocks in which trials were mixed but analysed separately for repeats of the same condition (e.g. visual reaction times quantified in trials following a visual trial) and switches between conditions (e.g. visual reaction times quantified in trials following an auditory trial). This revealed, for example, that the observed RSE was largest for unisensory blocks, hence when stimulus-response associations remained constant between trials, and was smallest for trials following a switch in sensory modalities (Gondan et al., 2004; Shaw et al., 2020). Further, this pinpoints a slowing of unisensory reaction times as underlying the difference in RSE between switch and repeat trials, while multisensory reaction times differed less between these types of trials. Hence, the relative slowing of unisensory reaction times following a switch of attention, cognitive resources or sensory-motor associations contributes to the observed multisensory RSE.

We here asked participants to detect supra-threshold visual and auditory stimuli in a classical RSE paradigm comprising visual, auditory, and audio-visual trials. Similar to previous work we presented these stimuli in three types of unisensory blocks and in mixed multisensory blocks, which comprised repeats and switches of the same modality. We then quantified the reaction times per sensory modality and condition (pure unisensory blocks, switches, repeats). As commonly done in multisensory studies, we also compared the observed multisensory reaction times to predictions by a race model, which predicts reaction times in dual-stimulus paradigms under the assumption of two independent processes and triggers a response whenever the faster of the two has finished (Raab, 1962). All in all, we found that task switching significantly contributes to the observed RSE and that violations of the race model emerge only in trials from mixed blocks but not when the three modalities are presented in isolation.

## 2. Material & methods

We tested 32 adult participants (mean age 24.5 years, SD 2.1, 9 males, 23 females). All provided a written informed consent before participating. Inclusion criteria were an age above 18 years, normal or corrected to normal vision, no self-reported hearing impairment, and no reported diagnosis of a neurological disorder. As compensation participants received 7€ per hour.

### 2.1 Stimuli and procedure

Experiments were performed in a darkened and sound-proof booth (E:Box; Desone, Germany). Participants sat comfortably in front of a monitor and keyboard and two speakers were positioned adjacent to the left and right of the monitor (Fig. 1A). Each trial started with the appearance of a centrally presented fixation cross, and after a random period (700 ms to 1100 ms uniform interval) one of three stimulus conditions (auditory, visual, or audio-visual) was presented. Inter-trial intervals lasted between 800 ms and 1200 ms (uniform). The auditory stimulus consisted of a pure tune (1000 Hz tone, 15 ms duration plus 5 ms rise/fall times, presented at 65 dB SPL over both speakers). The visual stimulus consisted of a white disk presented on a dark grey background for 16 ms (two refresh cycles), subtending 3 degrees of visual angle and was presented on a 27” monitor (ASUS PG279Q, 120 Hz refresh rate, gray background of 16 cd/m^2^). The audio-visual stimulus consisted of simultaneous presentation of the auditory and visual stimuli. Stimulus presentation was controlled using the Psychophysics Toolbox (Version 3.0.14; http://psychtoolbox.org/) using MATLAB (Version R2017a; The MathWorks, Inc., Natick, MA). The synchronization of the auditory and visual stimuli was confirmed using an oscilloscope.

**Figure 1.**
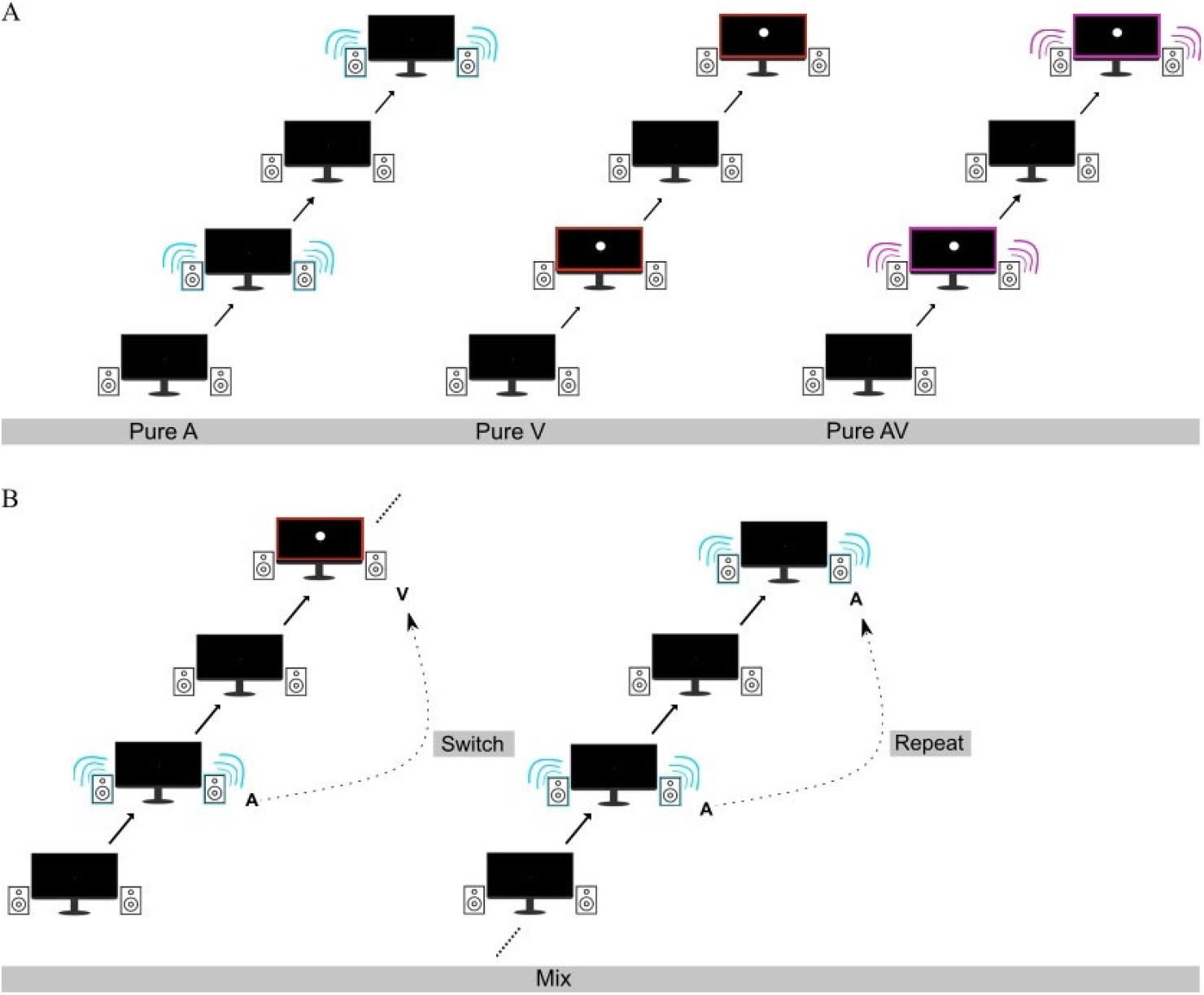
Experimental paradigm and conditions. Stimuli comprised either just an auditory stimulus (turquoise), just a visual stimulus (red) or an audio-visual stimulus pair (magenta), each preceded by a random fixation period. These were presented in pure unisensory blocks (**A**) or mixed blocks (**B**). For data analysis trials in the mixed blocks were divided into switches (A → V, V → A, A → AV, V → AV) or repeats (A → A, V → V, AV → AV).

Following the study by Shaw et al (Shaw et al., 2020) auditory (A), visual (V) and audio-visual (AV) stimuli were presented in different experimental contexts (blocks). First, we presented three types of unisensory blocks, containing either only auditory stimuli (**pure A**), only visual stimuli (**pure V**) or only audio-visual stimuli (**pure AV**). Second, there were **mixed** blocks, in which all three modalities were presented in a pseudo-random order. Participants task was to detect, as quickly as possible, the occurrence of either a visual or an auditory stimulus and to respond using a computer keyboard. Participants were allowed to use their preferred hand and fingers. Before the actual experiment, participants performed a test block of 20 trials, which consisted of the randomized presentation of all three modalities.

The focus of the study was to compare reaction times between pure blocks and trials from the mixed blocks that featured either a repeat of the same modality, or a switch between modalities (see definitions below). To ensure comparable trial numbers across modalities (A, V, AV) and conditions (pure, switch, repeat) we a priori created pseudo-random sequences of trials for the mixed blocks for each participant independently. The original design comprised 16 blocks per participant, 6 pure blocks (2 per modality) containing 100 trials each, and 12 mixed blocks containing 120 trials each. The sequence of mixed and pure blocks was pseudo-randomized for each participant prior to the study. Due to a mistake in the software running the experiment, only the first 12 blocks of this balanced and randomized design were presented in the experiment for each participant, and hence the effectively available number of trials per modality and condition differed. In the actual experiment, each participant performed 12 blocks, providing on average 790 trials (min 600, max 900) for the pure blocks, and on average 964 trials for mixed blocks (min 840, max 1094). Precise trial numbers available for data analysis are reported in Table 1.

**Table 1.** Trial numbers included in the analysis. Values reflect the median ([Min – Max]) per participant and analysis condition.

|  | Pure | Mix | Repeat | Switch |
| --- | --- | --- | --- | --- |
| Auditory | 271 [90 - 290] | 290 [234 - 335] | 142 [119 - 166] | 123 [90 - 142] |
| Visual | 253 [86 - 282] | 295 [246 - 342] | 139 [116 - 170] | 123 [100 - 149] |
| Audio-visual | 268 [85 - 287] | 289 [242 - 343] | 134 [115 - 167] | 130 [98 - 151] |

### 2.2 Data preparation and sorting into experimental conditions

The data were analysed offline in MATLAB 2019 using custom-made scripts and functions from the RaceModel Matlab package (https://github.com/mickcrosse/RaceModel). The data were pre-processed using by removing excessively fast or slow reaction times: we excluded the fastest and slowest 5% of trials for each participant across all blocks. Then, trials were sorted into the following conditions of interest (Shaw et al., 2020): Pure **A**, Pure **V** and Pure **AV** trials, obtained from the respective pure blocks. Then, we analysed **Mixed A**, **Mixed V** and **Mixed AV** trials, obtained from the mixed blocks regardless of the type of preceding trials. These mixed trials correspond to those usually investigated in classical RSE paradigms, where trials are analysed without considering the nature of the preceding trial. Finally, we split the trials from the mixed blocks into repeat and switch trials, as follows: **Repeat** trials were defined as those in which the stimulus in the analysed trial was the same as that in the preceding trial (i.e. **A** preceded by **A**, **V** preceded by **V**, **AV** preceded by **AV**). Switch trials were defined as those in which the stimulus in the analysed trial differed from that in the preceding trial (i.e. **V** preceded by **A**, **A** preceded by **V**, and **AV** preceded by **A** or **V**). As a result, the analysis of repeat and switch trials focused on distinct sets of trials from the mixed blocks, while the analysis of all mixed trials comprises both switch and repeat trials, and trials not included in either of these definitions (e.g. A or V trials preceded by AV trials).

### 2.3 Data analysis and statistics

Reaction times were analysed per sensory modality (A, V, AV) and condition (pure, mix, repeat and switch). In one analysis strand we computed for each participant, modality and condition the median reaction time (RT). These median RTs were then compared between modalities or conditions using paired Wilcoxon signed rank tests, corrected for multiple comparisons using the Bejnamini & Hochberg procedure for controlling the false discovery rate within the respective group of analyses. For each test we report the corrected p-values and the Z-value as a measure of effect size.

Following Shaw et al. (Shaw et al., 2020) we quantified the costs associated with switching modalities within the mixed blocks: these **switch costs** were obtained for each participant and modality as the median reaction time in switch minus that in repeat trials. We also quantified the costs associated with mixing tasks: these **mix costs** were obtained for each participant and modality as the median reaction time in repeat trials minus that in pure trials.

In a second analysis strand we compared the observed cumulative distribution functions (CDFs) of multisensory reaction times to a prediction under the assumption of two independent processes. More precisely, we used Raab’s model (Raab, 1962) to predict multisensory RTs under a probability summation account, in which on each trial the faster of two independent unisensory processes drives behaviour. If this model explains the data, the following inequality holds:

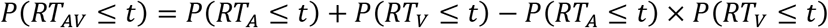

To obtain the model prediction, RT CDFs were evaluated at discrete quantiles (in 19 steps, between 5 and 95 in steps of 5) and the model prediction was calculated based on the observed auditory and visual CDFs using the OR model in the RaceModel package for Matlab. To quantify whetehr the observed data violate this model, we subtracted the prediction of this model (i.e. the right hand side) from the actually obserevd AV CDF. A positive difference is indicative of faster reaction times than expected under the probability summation account and is usually observed in multisensory detection tasks. To statistically compare the observed AV CDF with the model prediction, we performed separate one-sided paired Wilcoxon signed rank tests for each quantile and condition. We quantified the race model violation by calculating the positive area under the curve (AUC) of the difference, which provides an index of facilitative multisensory interaction (Miller, 1986; Nozawa et al., 1994; Shaw et al., 2020). We contrasted the obtained AUC values between conditions using paired Wilcoxon signed rank tests.

We implemented these analyses twice. Once using all trials for each participant and once after stratifying the number of trials across conditions and participants. This was done to ensure that differences in the available number of trials did not confound the results obtained from the full dataset. For this we used the same number of trials (n=89) per condition and participant, always selecting for each participant the first 89 trials from each condition and modality. This number was defined on the smallest number of trials across all participants and conditions.

## 3. Results

### 3.1 Median reaction times

In a first analysis, we compared reaction times (RTs) between modalities (A, V, AV) within each of the four conditions (pure, mixed, repeat and switch trials). As expected, this revealed that AV RTs were generally fastest (Fig. 2A). A group-level statistical analysis showed that AV RTs were significantly faster than the faster of the two unisensory modalities (A) for each of the four conditions (n=32 participants; paired Wilcoxon signed rank tests corrected using the Benjamini Hochberg procedure for controlling the false discovery rate; pure: p_corr_<0.0001, Z=4.73; mix: p_corr_ <0.0001, z=4.94; repeat: p_corr_ <0.0001, Z=4.83; switch: p_corr_ <0.0001, Z=4.94).

**Figure 2.**
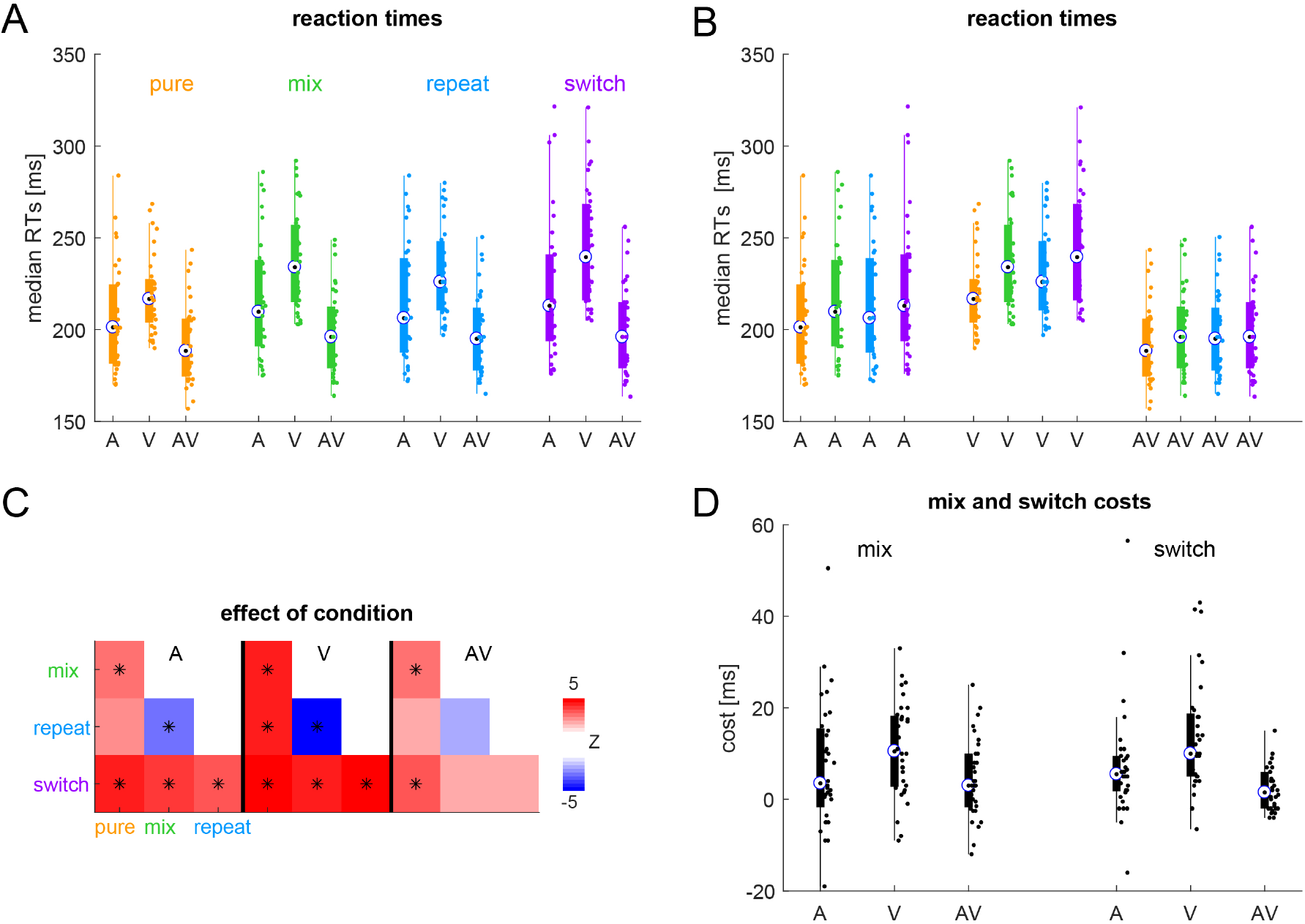
Reaction times. For each participant (dots) condition and modality we derived the median reaction time across trials. Boxplots show the group-median (circle) and the range of 25^th^ to 75^th^ percentiles across participants. **A.** Sorted per condition. **B.** Sorted per modality. **C.** Outcome of statistical contrasts between conditions, computed within each modality separately (paired Wilcoxon signed rank tests). P-values were corrected for multiple comparisons using the Benjamini Hochberg procedure for controlling the false discovery rate; * p<0.01). The color-code indicates the effect size (Z-value). **D.** Mixing and switching costs for each modality (see text for definition).

In a second analysis, we investigated the effect of condition separately for each modality (Fig. 2B). This revealed that pure RTs tended to be fastest and switch RTs to be slowest. This was statistically significant for the combinations indicated in Fig. 2C (paired Wilcoxon signed rank tests FDR corrected): in particular, for the unisensory conditions (A, V) switch trials were significantly slower compared to the other conditions (p_corr_ <0.01). For multisensory trials (AV) there were fewer significant differences (only pure vs. mix and switch trials, p_corr_ <0.01). To further probe whether the difference between unisensory and multisensory RTs variesd across conditions, we derived two types of costs describing the switching and mixing of modalities: first, the cost associated with switching sensory modalities (within the mixed blocks, defined as switch minus repeat trials), and the cost associated with mixing tasks (defined as repeat trials minus pure trials, hence separating the effect of block type) (Fig. 2D). Both costs were higher for unisensory compared to multisensory stimuli: statistical tests revealed significantly higher switch and mixing costs for visual compared to audio-visual stimuli (Mixing cost AV vs. A: p_corr_ =0.221, Z=1.23; AV vs. V: p_corr_ =0.02, Z=2.58; Switch cost AV vs. A: p_corr_ =0.029, Z=2.29; AV vs. V: p_corr_ <0.0001, Z=4.47). Hence, our data show that particularly the visual RTs were slowed by the mixing of sensory modalities within a block.

### 3.2 Race model violation

These results suggest that any benefit of multisensory over unisensory stimuli should be particularly pronounced in switch trials compared to pure trials. To confirm this, we contrasted the distributions of multisensory RTs with predictions from a race model, which provides an estimate of the expected RTs under the assumption of provability summation (Raab’s model, Raab, 1962). Figure 3A displays the measured (lines) and predicted (dots) cumulative RT distributions (CDFs’) for each condition and participant. Figure 3B directly visualizes the difference between the observed minus the model-predicted distribution; here, positive differences indicate a violation of the model’s assumption of two independent sensory channels driving behavior. Positive differences (model violations) are usually particularly prominent for faster RTs and are a prevalent test for a genuine benefit of multisensory information. We directly probed for statistical evidence for a violation of the race model, by contrasting the model prediction with the multisensory CDF across participants (one-sided paired Wilcoxon signed rank tests, FDR corrected across the 19 quantiles and conditions). This revealed the expected violations of the race model for conditions involving trials from mixed blocks (significant percentiles at p_corr_ <0.01 for mix trials 10% - 25%, maximal Z=3.9; repeat trials: 10% and 15%, maximal Z=3.4; switch trials: 10% to 35%, maximal Z=4.0), but not for the data from pure blocks (minimal p_corr_ was 0.12, maximal Z=1.89).

**Figure 3.**
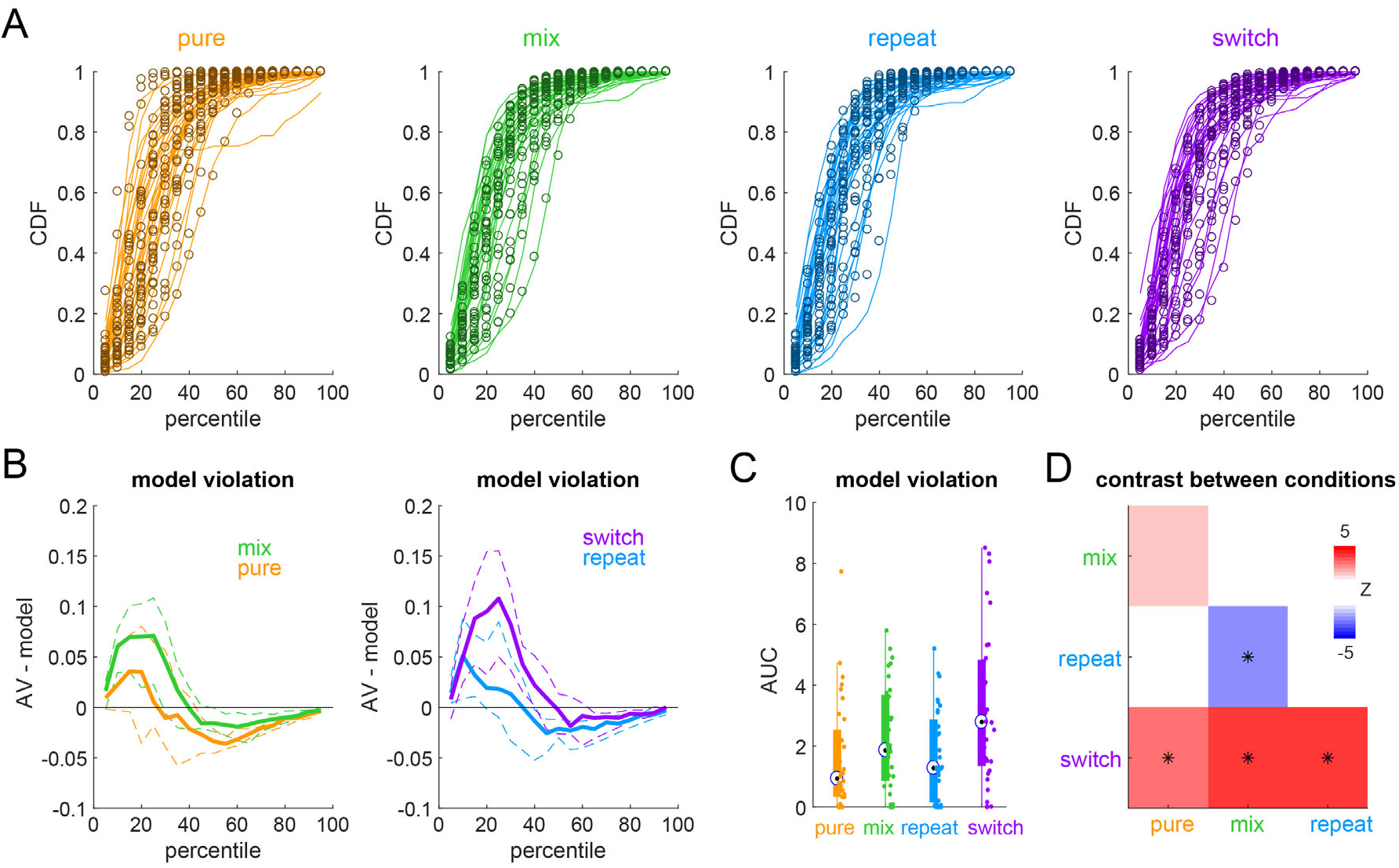
Race model analysis. **A.** Cumulative distributions (CDFs) of the reaction times in the audiovisual condition (lines) and the predictions from the Race model (circles) for each participant and condition. **B.** Violation of the race model, defined as the difference of the observed CDF in the audiovisual condition minus the model prediction. Thick lines indicate the group-median, dashed lines the two-sided 5% confidence intervals of the group-median derived from a percentile bootstrap distribution (2000 samples of participants). **C.** Violation of the Race model, defined as the (positive-) area under the curve (AUC) of the difference between true data minus model prediction. Boxplots show the group-median (circle) and the range of 25^th^ to 75^th^ percentiles across participants. **D.** Outcome of statistical tests contrasting the AUC values between conditions (paired Wilcoxon signed rank tests). P-values were FDR corrected; * p<0.01). The color-code indicates the effect size (Z-value).

Following previous work, we further quantified this violation using the area under the receiver operator characteristic (Fig. 3C). Model violation (AUC) was smallest for pure trials and largest for switch trials. A statistical analysis revealed that the AUC was significantly higher for switch trials compared to all other conditions (Fig. 3D, FDR corrected paired Wilcoxon signed rank tests; all p_corr_ <0.01), while it did not differ between the other conditions.

### 3.3 Control analyses

Due to a technical problem while running the experiment, the effectively available number of trials differed between participants, modalities, and conditions (Table 1). To confirm that our main results are not driven by this difference in the available number of trials we repeated the main analyses after stratifying the data to contain the same number of trials per participant, condition, and modality. In this stratified data, we still observed faster RTs in the AV condition compared to the faster unisensory condition (n=32 participants; paired Wilcoxon signed rank tests corrected using the Benjamini Hochberg procedure for controlling the false discovery rate; pure: p_corr_=0.0031, Z=2.96; mix: p_corr_ <0.0001, Z=4.91; repeat: p_corr_ <0.0001, Z=4.85; switch: p_corr_ <0.0001, Z=4.88). Mixing and switching costs also differed between unisensory and multisensory trials (mixing cost AV vs. A: p_corr_=0.0424, Z=2.15; V: p_corr_ =0.1471, Z=1.45; switch cost AV vs. A: p_corr_ =0.0389, Z=2.34; V: p_corr_ =0.0001, Z=4.18; FDR). Furthermore, the AUC model violation was again smallest for pure trials (median 0.53) and largest for switch trials (switch: 2.50, mix: 2.33, repeat: 1.48), and differed significantly between switch trials and the other conditions (switch vs. pure, mix and repeat: Z=2.71, Z=2.58 and Z=2.88, all p_corr_=0.019). We take this as evidence that differences in available number of trials do not drive the between-condition differences reported above.

## 4. Discussion

Reaction times for multisensory stimuli often appear faster than those for the respective unisensory stimuli. This multisensory redundant signal effect (RSE) has been replicated in many studies and has been considered as an important marker of a multisensory integration benefit (Miller, 1982; Schroger and Widmann, 1998; Minakata and Gondan, 2019). Typically, this RSE is experimentally obtained in paradigms in which the three experimental conditions are randomly mixed and hence the nature of the upcoming stimulus remains uncertain to the participant. As a result of this random mixing of conditions, the stimulus-response mapping that the brain employs for responding during each trial changes as well, an effect that resembles classical task switching (Rogers R. D. and Monsell, 1995; Wylie et al., 2003; Wylie et al., 2009; Sandhu and Dyson, 2013). Hence, the typical processing costs induced by task switching may potentially contribute to the experimentally measured speeding of multisensory responses.

### 4.1 Task switching slows unisensory reaction times in mixed blocks

The notion that task switching contributes to the measured multisensory RSE has been raised previously (Driver and Spence, 1998; Spence et al., 2001; Gondan et al., 2004; Otto and Mamassian, 2012; Shaw et al., 2020). Several studies suggested that the observed RSE depends on the order in which the three experimental conditions are presented. For example, Shaw et al. (2020) directly compared the RSE between an experimental design based on pure unisensory blocks, and a design employing mixed blocks but where data analysis separated trials featuring a repeat and those featuring a switch of the previous modality. This revealed that switching sensory modalities incurs switching and mixing costs known from typical cognitive tasks. These costs specifically affected the unisensory reaction times more than the multisensory ones and thereby expanded the numerically obtained RSE.

The present study was designed to directly replicate studies demonstrating that task switching costs conflate the multisensory RSE, based on the experimental design and analysis logic suggested previously (Gondan et al., 2004; Shaw et al., 2020). Our data confirm the contribution of task switching costs to the RSE. The numerical RSE was smallest in pure unisensory blocks, larger for trials repeating the same sensory modality within mixed blocks and was largest following a trial-by-trial switch in sensory modality. Furthermore, unisensory reaction times differed more between unisensory and mixed blocks than those to multisensory stimuli, confirming the hypothesis that the observed differences in RSE between pure, repeat or switch trials are a result of changes in unisensory perceptual-response processes rather than in genuine multisensory integration. Still, the precise causes of this slowing of unisensory reaction times remain unclear. In particular, whether this slowing rests on high-level cognitive control (Botvinick et al., 2001), processes involved in motor preparation (Giray and Ulrich, 1993), or potentially also results from low- and sensory-level processes such as attention (Miniussi et al., 1998; Schroger and Widmann, 1998; Talsma et al., 2010) remains to be determined.

### 4.2 The speeding of multisensory reaction times

For all trials from mixed blocks, the multisensory reaction times were significantly faster than unisensory reaction times and faster than those predicted by the race of two independent processes. Such a violation of the race model is often considered a hallmark of multisensory integration (Molholm et al., 2002; Gondan et al., 2004; Murray et al., 2005; Otto and Mamassian, 2012; Shaw et al., 2020). In particular, this violation has been taken to suggest that additional processes, such as a convergence of multisensory information prior to a decision process, must underlie behavior in multisensory conditions (Miller, 1982). Similar as the previous study by Shaw et al. (2020) we did not find a signification violation of the race model for reaction times in pure blocks. Hence, while multisensory reaction times were faster than unisensory ones in these pure blocks, there was no significant gain beyond that expected from redundant signals for switch trials, speaking against a significant and genuine multisensory benefit when task switching costs are experimentally reduced.

Previous work has remained controversial on whether a significant violation of the race model is seen when reaction times are obtained in pure unisensory blocks. While some studies did not find a violation of the race model (Crosse et al., 2015; Shaw et al., 2020), other studies using different designs or stimuli did find such a violation (Otto and Mamassian, 2012; Minakata and Gondan, 2019). Possibly the efficacy of the individual unisensory stimuli in eliciting rapid responses plays a central role for the expected benefit of multisensory stimuli (Minakata and Gondan, 2019). In addition, as highlighted by Otto and Mamassian (2012), the combined multisensory stimulus differs from the two separate unisensory stimuli not only by the combination of two sensory inputs, but also by an increased trial by trial variability of the underlying processes. This variability can contribute significantly to the observed deviation from a race model prediction. Systematic manipulations of stimulus salience or signal to noise ratio in future studies could help to directly test whether the observed RSE from pure unisensory blocks changes with the overall efficacy of the individual stimuli in driving behavior.

## 5. Conclusion

We probed to what degree task switching costs contribute to the apparent speeding of multisensory reaction times in paradigms intermingling multisensory and unisensory trials. Confirming previous studies, we observed significant task switching costs. In fact, when these were re experimentally reduced the multisensory redundant signals effect became negligible, suggesting that the benefit of multisensory signals for speeded detections may be less prominent than often upheld.

## Author contributions

CK: conceptualization, data analysis, writing and editing, supervision; MA, JH, JM, SÜ: data collection, data analysis, writing

## Data availability

The data file and Matlab code underlying the reported analyses are available from the bottom link on this resource page: https://www.uni-bielefeld.de/fakultaeten/biologie/forschung/arbeitsgruppen/cns/resources/index.xml Upon acceptance the data and code will be uploaded to a repository.

## Acknowledgements

This work was supported by funds from Bielefeld University.

